# Opposite effects of choice history and stimulus history resolve a paradox of sequential choice bias

**DOI:** 10.1101/2020.02.14.948919

**Authors:** Ella Bosch, Matthias Fritsche, Benedikt V. Ehinger, Floris P de Lange

**Affiliations:** Donders Institute for Brain, Cognition and Behaviour, Radboud University, Nijmegen, the Netherlands

**Keywords:** choice repetition, serial dependence, confidence, decision-making, adaptation

## Abstract

Perceptual decisions are biased towards previous decisions. Previous research suggests that this choice repetition bias is increased after previous decisions of high confidence, as inferred from response time measures (Urai et al., 2017), but also when previous decisions were based on weak sensory evidence (Akaishi et al., 2014). As weak sensory evidence is typically associated with low confidence, these previous findings appear conflicting. To resolve this conflict, we set out to investigate the effect of decision confidence on choice repetition more directly by measuring explicit confidence ratings in a motion coherence discrimination task. Moreover, we explored how choice and stimulus history jointly affect subsequent perceptual choices. We found that participants were more likely to repeat previous choices of high subjective confidence, as well as previous fast choices, confirming the boost of choice repetition with decision confidence. Furthermore, we discovered that current choices were biased away from the previous evidence direction, not previous choice, and that this effect grew with previous evidence strength. These findings point towards simultaneous biases of choice repetition, modulated by decision confidence, and adaptation, modulated by the strength of evidence, which bias current perceptual decisions in opposite directions.

## INTRODUCTION

Perceptual decisions are not only based on current sensory evidence, but also influenced by the choice history. Across a wide range of perceptual decision-making tasks, observers tend to repeat their decisions more than is expected by chance (Abrahamyan et al., 2016; Akaishi et al., 2014; Braun et al., 2018; Fischer & Whitney, 2014; Fritsche et al., 2017; Frund et al., 2014; St. John-Saaltink et al., 2016; Urai et al., 2017, 2019). This choice repetition bias does not only occur in human perceptual decision-making, but also in monkeys (Gold et al., 2008) and rodents (Busse et al., 2011; Odoemene et al., 2018), and has been found in other domains of human decision-making, such as free choice tasks and economic decisions (Allefeld et al., 2013; Padoa-Schioppa, 2013). In summary, choice history biases appear to be a general feature of decision-making.

Why does choice history bias occur? This is an especially good question given that, in most laboratory experiments, stimuli are uncorrelated across trials. Repeating previous choices is consequently detrimental to task performance; choice history biases are maladaptive in the contexts of such tasks. However, they are likely adaptive in natural conditions, as our environment usually remains relatively stable over short timescales (Dong & Atick, 1995; Simoncelli & Olshausen, 2001). Crucially, observers can exploit this stability by leveraging information from the recent past in order to stabilize perceptual decisions against disturbing factors, such as noise (Cicchini et al., 2017; Fischer & Whitney, 2014). Temporally smoothing internal representations in this manner would manifest itself in a tendency to repeat previous decisions. Choice history biases that persist despite uncorrelated input may be a consequence of our prior expectation that our environment tends to be temporally correlated.

Bayesian theories of perceptual decision-making prescribe how previous information should be integrated with current information in a probabilistically optimal manner (Vilares & Kording, 2011). Such theories would predict that current choices should be more strongly biased to previous choices when the previous choice is associated with high certainty. In line with this idea, several studies have found that choice repetition is stronger when the previous choice was fast and when arousal was low (Braun et al., 2018; Urai et al., 2017) - two factors which have been linked to increased decision confidence (Sanders et al., 2016; Urai et al., 2017). Moreover, recent studies using a continuous estimation tasks found that a higher self-reported decision confidence on the previous trial was associated with a stronger bias on the current trial towards the previous perceptual estimate (Samaha et al., 2019; Suárez-Pinilla et al., 2018). Thus, broadly in line with Bayesian theories, it appears that high decision confidence on the previous trial leads to a stronger choice repetition bias.

Surprisingly however, it has also been reported that observers are more likely to repeat a previous choice that was based on low, compared to high, sensory evidence (Akaishi et al., 2014). According to Akaishi and colleagues, choice repetition arises from internal signals as previous choices shift the internal choice estimate, biasing the subsequent choice. They argued that the estimate updates more towards a choice made based on low sensory evidence.

Crucially, there is an apparent contradiction: Akaishi finds that choice repetition is strongest when the previous choice was associated with *low* confidence (indicated by low sensory evidence), whereas Urai and Brain found that choice repetition is strongest when the previous choice was associated with *high* confidence (indicated by fast responses and low arousal). This paradoxical state of affairs suggests that there are several, possibly interacting, factors that jointly determine the presence and strength of serial choice biases.

We set out to isolate the effects of choice history and stimulus history on serial choice bias by examining how different factors modulate choice repetition probability. Participants performed a motion coherence discrimination task, identifying test stimuli as either more or less coherent than a reference stimulus while also reporting their subjective decision confidence. Stimulus evidence was parametrically varied using 6 levels of evidence strength. This allowed us to examine the effect of previous decision speed and previous decision confidence, as well as previous stimulus evidence, on choice repetition.

We find that choices are biased towards the previous choice, and that this bias is stronger for confident as well as fast previous choices. This is in line with previous findings using response times and pupil dilation as proxy measures for confidence (Braun et al., 2018; Urai et al., 2017). In addition, we find that choices are biased away from the direction of evidence on the previous trial, and more so when the evidence was strong, explaining the findings of Akaishi et al. (2014). Taken together, perceptual choices are biased towards the previous choice, a modulation that grows with previous decision confidence, and biased away from the previous evidence direction, a modulation that grows with previous evidence strength. These findings suggest that previous choices and previous stimuli may induce biases on separate stages of perceptual decision-making.

## METHODS

### Data availability

All data and code used for stimulus presentation and analysis will be made available from the Donders Institute for Brain, Cognition and Behavior repository at https://data.donders.ru.nl/collections/di/dccn/DSC_3018029.09_654?5.

### Participants

Thirty-eight naïve participants (23 female/15 male, age range 18-34 years) recruited through the university pool took part in the experiment. Subjects were paid 8 euros an hour for their participation. All participants reported normal or corrected-to-normal vision and provided written informed consent before the start of the study. The study was approved by the local ethical committee (CMO region Arnhem-Nijmegen, The Netherlands) and was in accordance with the Declaration of Helsinki (2008 version).

We performed an a-priori power analysis that resulted in *n* = 34 to obtain 80% power for detecting at least a medium effect size (*d* ≥ 0.5) with a two-sided paired t-test at an alpha level of 0.05. Four participants were excluded from our original sample: one did not complete all sessions, one was excluded after training due to failure to follow task instructions, and two were excluded due to technical errors during the experiment. These participants were replaced with new participants.

### Apparatus & stimuli

Visual stimuli were generated with the Psychophysics Toolbox (Brainard, 1997; Kleiner et al., 2007; Pelli, 1997) for MATLAB (2018). They were displayed on a 24″ flat panel display (Benq XL2420T, resolution 1920 × 1080, refresh rate: 60 Hz). Participants viewed the stimuli from a distance of approximately 70 cm in a dimly lit room.

All stimuli were random dot kinematograms composed of 769 white dots on a black screen, moving within a central circular aperture (12 deg visual angle radius). The dot density was 1.7 dots per deg^2^. A red fixation cross was displayed at the center of the screen at all times. The population of dots was split into “signal dots” and “noise dots”. The signal dots moved in the motion direction of the trial with a velocity of 11.5°/s. If signals dots left the aperture, they were redrawn on the opposite side. Three different sequences of dot motion (at the same coherence and direction) were presented in an interleaved fashion, making the effective speed of signal dots 3.83°/s. The noise dots changed position randomly from frame to frame. The percentage of signal dots defined the motion coherence, a measure of motion strength.

### Procedure

In each trial of the experiment, two white random dot motion stimuli were presented on a black background successively for 750 ms, separated by a 250 ms inter-stimulus interval. The first stimulus was always a reference stimulus of 70% motion coherence. The second stimulus was a test stimulus with a higher or lower motion coherence than the reference. The difference in motion coherence between reference and test stimuli was taken from one of three sets, chosen on a participant-by-participant basis (procedure described below): easy (1.25, 2.5, 5, 10, 20 and 30%), medium (0.625, 1.25, 2.5, 5, 10 and 30%) or hard (0.3125, 0.625, 1.25, 2.5, 5 and 20%). Both stimuli had the same mean motion direction, and the motion direction of any given trial was randomly offset between 30° and 330° from the motion direction of the previous trial.

Participants were asked to give two responses in order: first, they indicated whether the test stimulus had lower or higher coherence than the reference (coherence response); second, they reported how confident they were about their decision on a 1-4 point scale (confidence report). For the coherence response, they used their right hand to press either the ‘j’ or ‘k’ button on the keyboard. The button mapping for indicating lower or higher coherence of the test stimulus was counterbalanced across participants. For the confidence report, participants used their left hand to press the corresponding 1-4 digit buttons on the left-hand side of the keyboard. Participants had 4.75 seconds to give both responses, starting from the onset of the test stimulus. If they failed to give both responses in the correct order within the time limit, they received auditory feedback consisting of a low tone, played through headphones during the inter-trial interval. The sequence of coherence differences between the reference and test stimuli was pseudo-randomized across trials, such that every coherence difference was preceded equally often by every other coherence difference (Brooks, 2012).

Participants completed three sessions of which one practice session and two data collection sessions. During the practice session, participants received instructions about the coherence discrimination task and performed one or more simplified practice blocks of 48 trials each, in which they only had to judge the coherence difference without rating their confidence. Next, a staircasing procedure was used to estimate an individual threshold of 70% accuracy in the coherence discrimination task using the QUEST algorithm (Watson & Pelli, 1983). Participants completed at least 3 blocks of 48 trials each after each of which the convergence of the threshold estimate was visually inspected. Based on the resulting threshold, one of the three stimulus sets was chosen: for thresholds below 5% and below 10% coherence difference, the hard and medium stimulus sets were selected, respectively. As a result, 2 participants were assigned the easy set, 22 participants the medium set, and 10 participants the hard set. After the staircasing procedure, participants received instructions for the additional confidence report and practiced the complete task with their stimulus set for the rest of the first session (9 blocks of 48 trials, 432 trials total).

The two data collection sessions started with one refresher block of 48 trials. Then participants completed 15 main blocks of 48 trials for each session, resulting in 1440 total trials per participant.

Participants received auditory feedback about the correctness of their decision during the practice blocks and refresher blocks only. This feedback consisted of a brief high or low tone for correct and incorrect decisions, respectively, played through headphones during the inter-trial interval. Participants always received on-screen written feedback about their general performance (percentage correct, average response time, and missed trials) in each block.

### Data cleaning

Trials in which one or both responses were missing and trials where participant gave a coherence response within ≤ 300 ms from the onset of the test stimulus were removed from further analyses. Consequently, 184 out of 48,960 trials (0.38% of all trials) were discarded.

### Deriving choice repetition from psychometric functions

In order to quantify the choice repetition bias and its modulation by previous evidence, response time, and confidence, and qualitatively compare findings to previous studies, we first employed a psychometric function fitting approach. We estimated choice repetition independently for each condition, following the analytical approach of earlier studies (Akaishi et al., 2014; Urai et al., 2017).

We first expressed the probability of a “higher coherence” response (*P*(*r*_*t*_ = 1)) as a function of the signed coherence difference between the reference and test stimulus 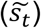 and fit a psychometric function (Figure 2a; Wichmann & Hill, 2001) of the form

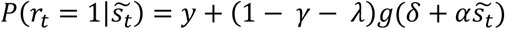

where *λ* and *γ* were the probabilities of stimulus-independent errors (‘lapses’), *g* was the logistic function, *α* was perceptual sensitivity, and *δ* was a bias term. The free parameters *λ, γ, α* and *δ* were estimated by minimizing the negative log-likelihood of the data (using MATLAB’s *fminsearchbnd*). We constrained *λ* and *γ* to be identical to estimate a single, choice-independent lapse rate.

**Figure 1:**
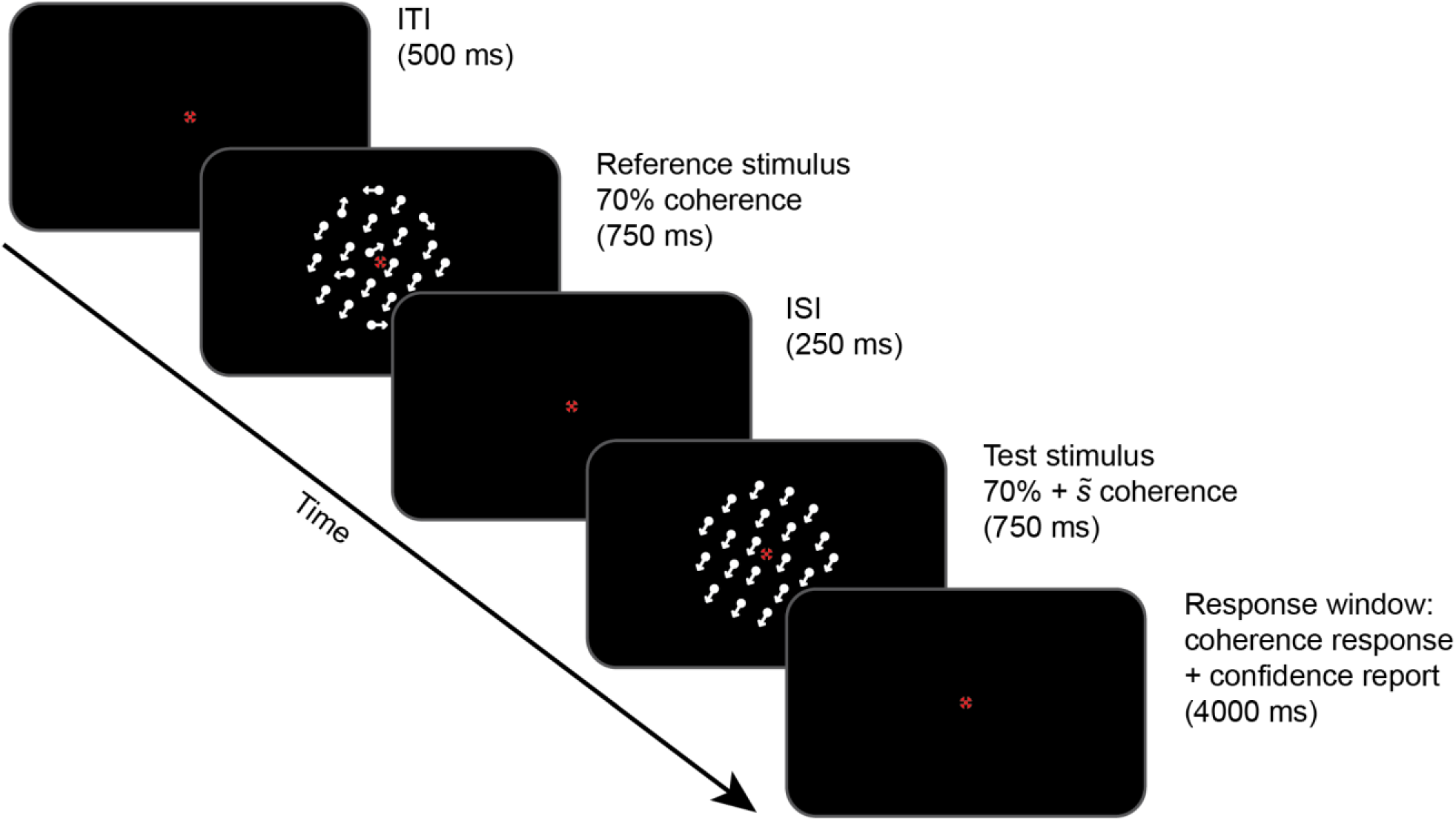
Trial design of main task. A reference random dot motion stimulus of 70% coherence was presented at fixation, followed by a test stimulus with a different coherence but with the same mean motion direction. Participants gave two responses: first, they indicated whether the test stimulus had higher or lower coherence than the reference, using the ‘j’ and ‘k’ button on the keyboard; second, they reported their confidence on a scale of 1-4. If they failed to give both responses, they received auditory feedback during the ITI.

**Figure 2:**
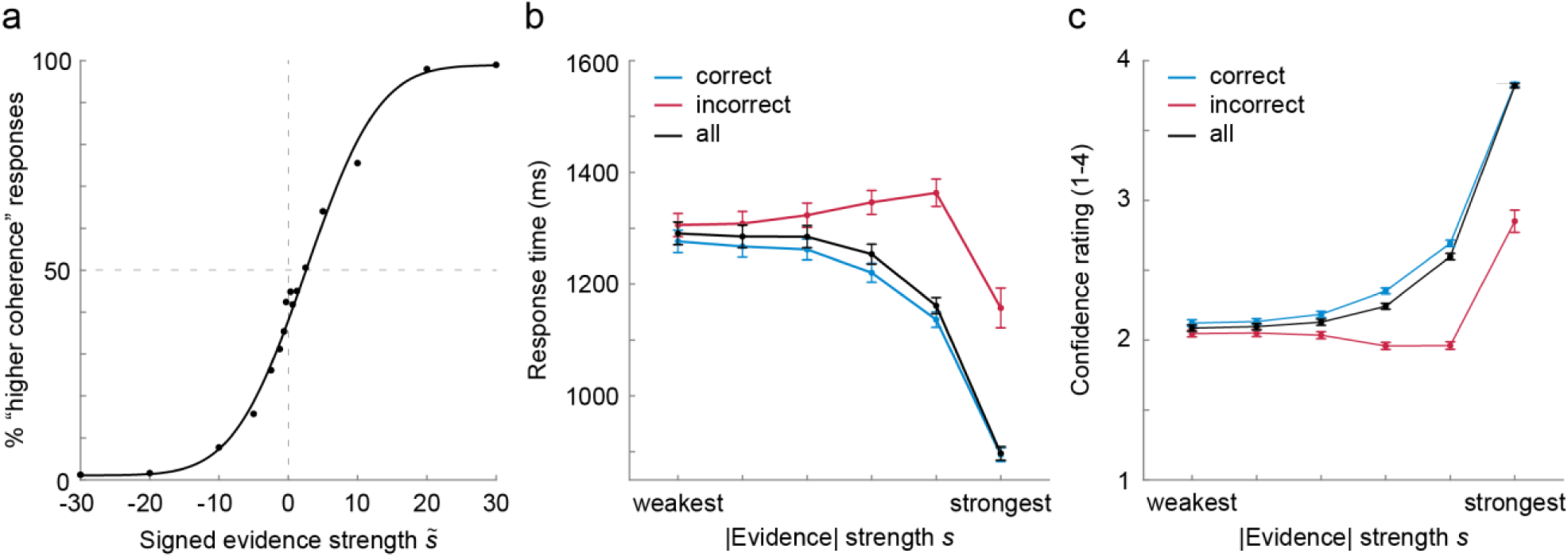
Coherence discrimination performance. (a) Group average responses follow a psychometric function, with a general bias towards “lower coherence” responses. (b) Mean reaction times decrease with absolute evidence for correct choices and increase for incorrect choices, except for trials with the largest sensory evidence. (c) Mean subjective confidence ratings increase with absolute evidence and decrease for incorrect choices, except for trials with the largest sensory evidence. Error bars represent between subject *SEM*s.

For the quantification of serial choice bias, we first split the data into two bins corresponding to the previous choice such that one bin contained all trials for which the participant previously reported “higher coherence” while the other bin contained all trials for which they reported “lower coherence”. For each level of previous absolute evidence strength (*s*_t-1_) within these bins, we further split the data by previous response time (*rt*; based on a median-split) or previous confidence (low ratings (1, 2) and high ratings (3, 4)). For each of those subsets of trials, we fit the psychometric function as described above. In order to compute the choice repetition bias, the resulting bias terms *δ* were transformed from log-odds into probabilities by the inverse logit function *P* = *e*^*δ*^/(1 + *e*^*δ*^). This probability reflects 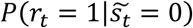, which is the probability to choose “higher coherence” in a hypothetical ambiguous trial (no evidence) in the current trial. For each bin we subtracted these values across the two previous choice options, which yielded the pooled measure of choice repetition probability. Finally, to test for differences in choice repetition probability, or p(repeat), across bins, we performed repeated measures ANOVAs using SPSS (*IBM SPSS Statistics for Windows*, 2015).

### History-dependent multiple regression model

While the approach described above allowed us to compare the current results to previous studies on choice repetition, it suffers from the problem that previous trial characteristics, such as evidence strength, response time, and confidence report are correlated (Figure 2). Splitting data according to one of these variables will partition meaningful variance in the other variables as well, which can introduce or mask apparent influences of any one variable on choice repetition. Furthermore, since the above analysis is focused on the biasing influence of the previous choice, it is not clear what role the previous stimulus information itself plays in biasing subsequent visual processing. To overcome these problems, we devised a history-dependent regression model, which allowed us to estimate separate influences of current and previous stimulus variables and response variables.

Specifically, we constructed a generalized linear mixed model (GLMM) with a binomial link function to predict the current choice based on current and previous stimulus and response variables as well as their interactions. The factors in this regression model can be conceptually split into current-trial factors and history factors, where the current-trial factors describe the stimulus information (i.e. evidence direction, evidence strength, and interactions) on the current trial, and the history factors describe the stimulus information and response characteristics (i.e. choice, response time, confidence report, and interactions) of the previous trial.

We were interested in the influence of the previous choice on current choice. Accordingly, we added the effect of previous choice (*prev choice*) as a factor to the model. To examine whether the influence of the previous choice is larger when participants were confident about that choice, we included an interaction factor (p*rev choice x prev confidence*) to the model. Similarly, to examine whether the influence of the previous choice was greater when participants had responded quickly, we also added this interaction factor (*prev choice x prev rt*). Furthermore, as the influence of the previous choice may scale with the strength of absolute evidence for that choice (i.e. the coherence difference between the reference stimulus and test stimulus), we also included this interaction factor (*prev choice x prev* |*evidence*|). Note that these three interactions are all theoretically related to decision confidence: more evidence leads to a more confident decision, just as a lower response time and higher reported confidence reflect a more confident decision.

It is important to note that, due to the difficulty level being staircased, there is a ∼70% correlation of previous choice and previous evidence direction (i.e. the sign of the evidence, determining whether it was a “higher coherence” or “lower coherence” trial). This raises the question whether it is the previous choice or the previous evidence direction that influences the current perceptual decision. To investigate this, we added the previous evidence direction (*prev evidence dir*) to the model, as well as all interactions equivalent to those we included for previous choice. This included interactions with previous confidence (*prev evidence dir x prev confidence*), previous response time (*prev evidence dir x prev rt*), and previous absolute evidence (*prev evidence dir x prev* |*evidence*|, equivalent to the signed evidence of the previous trial).

All factors included thus far describe history effects. However, observers’ decisions are primarily based on the bottom-up information present in the current trial. To account for this, we included the signed evidence of the current trial to the model (*curr evidence dir x curr* |*evidence*|).

Finally, we included the main effects of all variables in the aforementioned interactions (with the exception of *prev choice* and *prev evidence dir*, which were already included). Accordingly, we included *prev confidence, prev rt, prev evidence*, and *curr evidence* as factors to the model. Note that these main effects by themselves provide no information about the identity of either the previous or current trial, nor information about the previous choice, and were therefore unlikely to provide information about current choice. Consequently, they were not expected to be significant factors in the model. The reason they were nevertheless added was to prevent that an unexpected significant modulation would express itself as an interaction and hence be misinterpreted.

Before constructing the model, variables were re-coded as follows. Categorical predictors *choice* and *evidence dir* were coded using effect coding (−1/1). Confidence was subject-wise centered and subject-wise scaled by its standard deviations. For the response times we used a robust z-score and removed the subject-wise median and scaled by the subject-wise median absolute deviation (constant = 1.48). We scaled the unsigned evidence to range between 0 and 3, to accommodate smaller parameter estimates to prevent numerical floating-number overflow.

We used the R-package lme4 (Bates et al., 2015) to fit a generalized linear model from the binomial family. We fitted a model with ‘subjects’ as the only random grouping factor. We included for each fixed effect its corresponding random slope coefficient, but without random correlations, as the model did not converge. Even with this simplification, the random effect structure was singular, but the model converged according to the lme4 convergence checks. As a robustness check, we re-fit the data with a Bayesian GLMM using *brms* (Bürkner, 2017, 2018) with an LKJ-prior of 2 on the correlation matrix which confirmed all of our findings. For significance testing we report Walds-Z test. Walds Z-test is valid only in the asymptotic regime assuming a multivariate normal sampling distribution of parameters and a proportional sampling distribution of the log likelihood to *χ*^2^. Therefore, we will be very conservative in our interpretation of the reported p-values if the effects are not obvious from effect-sizes alone. To check whether the model can adequately capture our data, we plotted and compared fitted marginal against aggregated raw marginal data. Mimicking posterior predictive tests, we simulated new datasets from our model and compared the observed simulated data distributions with the observed one and generally found our data to be well captured.

## RESULTS

The goal of the current study was to investigate the modulation of sequential choice biases by subjective decision confidence, motivated by the seemingly conflicting roles of previous response times and stimulus evidence.

To this end, thirty-four human observers performed a binary forced choice coherence discrimination task on random dot motion stimuli. As expected, stronger absolute evidence resulted in higher accuracy (Figure 2a), faster response times (Figure 2b), and higher subjective confidence reports (Figure 2c). In addition, higher confidence reports were associated with higher accuracy and faster response times, which are often considered an implicit measure of decision confidence. These findings suggest that the subjective confidence reports are a meaningful reflection of decision confidence.

Both response times and confidence reports exhibited patterns corresponding to decision uncertainty (Sanders et al., 2016). Response times decreased with evidence for correct responses and increased with evidence strength for incorrect responses (Figure 2b), whereas confidence reports increased with evidence strength for correct responses and decreased with evidence for incorrect responses (Figure 2c). An exception to this pattern were incorrect responses based on the strongest evidence, where response times were faster and confidence reports higher.

### Choice repetition increases after fast choices

In line with previous research (Urai et al., 2017), we observed an increase of choice repetition after fast choices (Figure 3b). We confirmed these observed patterns by performing a repeated measures ANOVA, testing the effect of previous absolute evidence strength and previous response time on the choice repetition values we derived. This revealed a main effect of previous response time, with increased choice repetition for previous fast compared to slow choices (main effect of previous response time: *F*(1,33) = 32.708, *p* < .001). In addition, we found an interaction between previous response time and previous evidence (*F*(5,165) = 2.961, *p* = .014), indicating that the effect of previous response time was not equal across all levels of previous evidence (see Figure 3b). This main effect of response time was confirmed by our history-dependent regression model, indicating that previous response times negatively modulated the impact of the previous on the current choice (Figure 4; *prev choice x prev rt*: *b* = - 0.12; bootstrapped 95% *CI*s = −0.14, −0.09; *p* < .001). In other words, participants were more likely to repeat their choice after a fast response.

**Figure 3:**
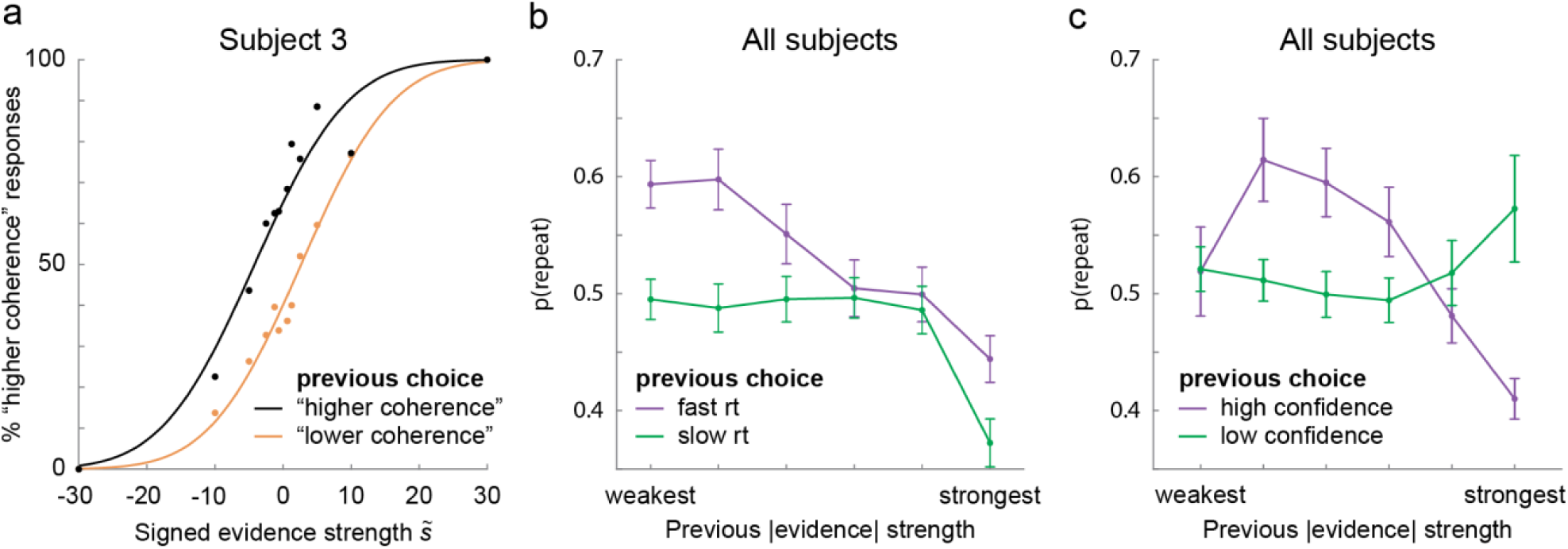
Replication of previous findings of choice repetition and variables that modulate choice repetition: previous evidence, previous *rt*, and previous confidence. P(repeat) values, reflecting choice repetition probability, are based on psychometric curve fitting after binning of the data. (a) Responses of an example participant, split for previous choice, reveal a clear choice history bias in this participant. The difference (δ) between the curves at 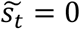 can be converted into a p(repeat) value using the inverse logit function. (b) Group p(repeat) values for previous fast response times versus previous slow response times (median split per evidence bin) show that choice repetition is higher after fast responses. Paradoxically, choice repetition decreases with previous absolute evidence. (c) Group p(repeat) values for previous highly confident versus previous low confidence show a varying modulation of choice repetition by previous confidence. Error bars represent between subject *SEM*s.

**Figure 4:**
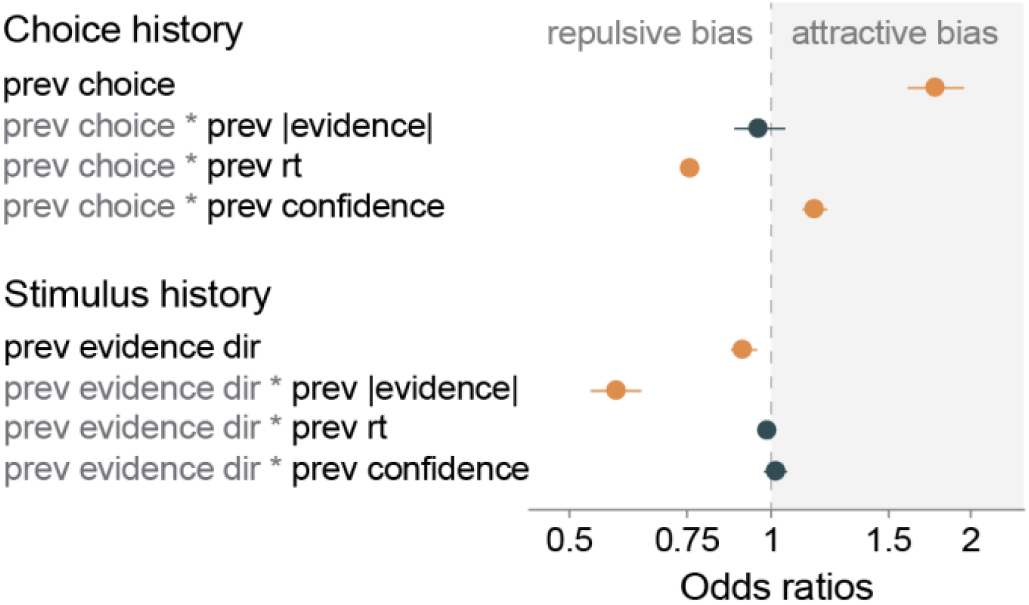
GLMM fixed effects. Odds ratios <1 signify a negative estimate, meaning the higher the term, the lower the current choice (and thus the more likely participants answered “less coherent”; odds ratios >1 imply a positive estimate. Significant terms (*p* < 0.05) are marked in orange. Error bars indicate *CI*s.

### Choice repetition increases after confident choices

This modulation by response time has been previously interpreted as evidence that choice repetition increases after confident responses. We next sought to test the role of confidence more directly by relating choice repetition to explicit subjective confidence ratings. To this end, we performed a repeated measures ANOVA on the pattern observed in Figure 3c, testing the effect of previous confidence rating and previous absolute evidence strength on the choice repetition values we derived. This showed a complexly varying modulation of choice repetition by previous subjective confidence across different levels of previous evidence strength (Figure 3c; interaction previous confidence and previous evidence strength: *F*(5,165) = 7.918, *p* < .001; main effect of previous confidence: *F*(1,33) = .444, *p* = .510). It is likely that the effect of previous confidence was difficult to derive from this analysis because of multicollinearities in the data. The history-dependent regression model accounts for this, and clearly revealed that previous confidence positively modulated the impact of the previous on the current choice (Figure 4; *prev choice x prev confidence*: *b* = 0.066; bootstrapped 95% *CI*s = 0.03, 0.10; *p* < .001). In other words, participants were more likely to repeat previous choices made with high confidence, even after adjusting for previous response time and previous evidence strength. We found that choice repetition increased following high subjective confidence reports.

### Choice alternation after previous high stimulus evidence resembles adaptation

Next, we investigated whether choice repetition was modulated by previous evidence strength. In line with Akaishi et al. (2014), the psychometric analysis showed that choice repetition decreased after stronger previous evidence strength (Figure 3b; main effect of previous evidence *F*(5,165) = 13.550, *p* < .001). This may appear paradoxical in light of our earlier described findings, as trials with strong evidence have faster response times and higher confidence ratings (Figure 2), which would be expected to lead to an increase in choice repetition. Strikingly, our model revealed that choice repetition was not significantly modulated by the strength of the previous evidence (*prev choice x prev* |*evidence*|: *b* = −0.017; bootstrapped 95% *CI*s = −0.10, 0.05; *p* = .662), contradicting our previous psychometric analysis. These contradicting findings raise the question if, and how, the strength of the previous evidence modulates the current choice.

The answer to this question may lie not with the choice history, but with the stimulus history. The previous stimulus information could potentially affect the encoding of current stimulus information, which in turn could bias the current choice (Kohn, 2007; Thompson & Burr, 2009; M. A. Webster, 2012, 2015). It is therefore important to account for the influence of the previous stimulus information.

Indeed we found that whereas current choices are biased towards the previous choice (*prev choice*: *b* = 0.25; bootstrapped 95% *CI*s = 0.16, 0.35; *p* < .001), they are simultaneously biased away from the previous evidence direction (*prev evidence dir*: *b* = −0.041; bootstrapped 95% *Ci*s = −0.08, 0; *p* = .042). This is particularly noteworthy as evidence direction and choice were correlated: 70.4% (*SD* = 3.8% across subjects) of choices corresponded to the evidence direction. Yet, the coefficients have opposing signs. We also found that the repulsion away from the previous evidence direction increased when this evidence was stronger (*prev evidence dir x prev* |*evidence*|: *b* = −0.23; bootstrapped 95% *CI*s = −0.30, −0.15; *p* < .001). This resembles a classical adaptation effect, which is typically stronger following strong adaptor stimuli (Thompson & Burr, 2009). However, note that in our design, observers’ choices are based on the coherence difference between two dot motion stimuli, and it is this difference that induces the observed effect. Consequently, our observed effect is not identical to a classical adaptation effect.

These findings reveal that the apparent and puzzling increase of choice repetition with decreasing evidence strength found by Akaishi et al. (2014) (Figure 3b) may in fact not be a modulation of the influence of the previous choice, but a modulation of the influence of previous sensory evidence on current sensory processing. In summary, we find that choices are biased towards the previous choice, a bias that grows when people were faster and more confident on the previous trial. Simultaneously, choices are biased away from the previous evidence direction, a bias that grows for stronger previous evidence (Figure 4).

## DISCUSSION

Perceptual choices are not only based on current sensory information, but are also systematically biased towards the recent history of previous choices. In the current study, we set out to test how choice repetition bias is modulated by aspects of the choice history as well as stimulus history. Specifically, we investigated the role of decision confidence on the probability to repeat the same choice on successive trials. Confidence deduced from response times and pupil dilation suggest that people are more likely to repeat previous choices (Braun et al., 2018; Urai et al., 2017), in line with an optimal integration of previous and current information in a stable environment. In apparent conflict, confidence deduced from sensory evidence suggests that observers are more likely to alternate from previous confident choices (Akaishi et al., 2014). To resolve this conflict, we measured decision confidence with explicit confidence ratings where previous studies probed decision confidence indirectly via response times or pupil dilation.

We found that observers were more likely to repeat confident as well as fast choices. This is in line with the previous findings from indirect measures of decision confidence and confirms the role of decision confidence in positively modulating choice repetition. Furthermore, we found that choice repetition decreased with increasing evidence strength on the previous trial, in line with Akaishi et al. (2014). Crucially however, our history-dependent regression model revealed that previous evidence did not modulate the transfer of successive choices, but rather the influence of the previous trial, as current choices were found to be biased away from the evidence direction on this previous trial.

### The role of decision confidence on choice repetition

According to Bayesian theories of perceptual decision-making, prior information is integrated with sensory input in a probabilistically optimal manner (Ernst & Banks, 2002; Vilares & Kording, 2011). Such theories would predict that prior information is leveraged more strongly if the uncertainty associated with this information is low; consequently, perceptual choices should be more strongly biased towards previous confident choices. Our findings are in line with these predictions: people are more likely to repeat previous confident choices. However, we cannot conclude from our data that this integration occurs as described by, and is optimal according to, Bayesian theories.

An open question is what underlying mechanism is modulated by decision confidence. One candidate mechanism would be the process of evidence accumulation, as recent finding shows that choice history biases are explained by a history-dependent change in the evidence accumulation (Urai et al., 2019). Confidence could modulate the influence that the previous choice exerts on the slope of evidence accumulation during the formation of subsequent choices. Specifically, confidence could make the slope steeper, thus increasing the likelihood of choice repetition.

Our findings add to the literature describing choice history effects of series of forced choices between two alternatives. However, most perceptual decisions are not binary, but continuous. In continuous estimation tasks, choices are serially dependent as observers are biased towards previous choices (Fischer & Whitney, 2014; Fritsche et al., 2017; Samaha et al., 2019; Suárez-Pinilla et al., 2018). It has been shown that this serial dependence biases increase following high subjective confidence reports (Samaha et al., 2019; Suárez-Pinilla et al., 2018). In line with these findings, when decoding uncertainty from early sensory areas, behavioral serial dependence is stronger when going from low to high levels of uncertainty (the inverse of confidence) compared to vice versa (van Bergen & Jehee, 2019). Our findings suggest a similar influence of confidence in forced choice paradigms. However, it is unclear to what extent choice repetition in forced choices and serial dependence in continuous estimations rely on the same underlying processes. More research is needed to synthesize findings from choice repetition and serial dependence in estimation tasks.

Previous studies into the influence of confidence on choice repetition in forced choice paradigms used proxy measures for confidence, such as pupil dilation and response times. In this study, we assess confidence using a subjective report measure and also measure response times, and find that both modulate choice repetition. An open question that remains is whether subjective confidence and confidence as assessed with response times have independent contributions to choice repetition probability. It should be noted that in our data, confidence reports and response times were correlated with each other and were both correlated with evidence strength, which modulated the influence of the previous stimulus on the current choice. One may wonder whether this multicollinearity in our data affects the interpretability of our model estimates. However, the reported confidence intervals of the parameter estimates suggest that the influence of these variables is robust, which points towards the interpretation that all these variables have independent contributions to choice repetition modulation. Further research needs to be conducted to further investigate this interpretation.

### The role of stimulus history on choice repetition

At first glance, our finding that observers’ choices are biased away from the previous evidence direction and that this bias grows with evidence strength resembles a sensory adaptation effect as frequently described in the literature (Kohn, 2007; Thompson & Burr, 2009; M. A. Webster, 2004, 2012, 2015). However, it is important to consider that in our experimental design, observers made judgments about the *difference* between a reference stimulus and a test stimulus.

Our history effects cannot be explained by sensory adaptation to motion. The main reason for this is that we changed the motion direction at least 30 degrees from one trial to the next. The varying motion direction should prevent any patterns of motion adaptation across trials. Sensory adaptation can however explain the within-trial effect of a general response bias towards perceiving the test stimulus as less coherent than the reference stimulus (Figure 2a), as within a trial both stimuli always had the same motion direction.

As our findings resemble but cannot be explained by sensory adaptation, this raises the question from which neural population(s) and at what stage of the decision formation process our adaptation-like modulation arises. We hypothesize that the adaptation arises not from a population of sensory neurons which encode the motion coherence, but rather from a neural population which encodes the accumulated (difference in) motion coherence. Also, we hypothesize that neural adaptation away from this difference should occur at a stage after this difference is computed from the bottom-up sensory evidence but before this evidence is converted into the final choice. After all, an adaptation to the choice itself would predict choice alternation, whereas we find choice repetition. Follow-up research should investigate these hypotheses about the neural and temporal characteristics of the adaptation-like modulation we describe.

Some previous research has described both attractive choice biases and repulsive stimulus biases in perceptual decision-making. Fritsche et al. (2017) found in seperate orientation estimation tasks that observers’ perception of stimuli was repulsed by previous stimuli while their choices were attracted towards previous choices. More recently, Fornaciai & Park (2019) showed a repulsive adaptation effect in the absence of an attractive serial dependence effect, when they removed the influence of late modulatory feedback by visual backward masking. These findings suggested that attractive and repulsive effects may jointly but independently contribute to perceptual experience. Indeed, we show that choice history and stimulus history bias our perceptual experience in opposite directions yet in tandem.

## CONCLUSIONS

We find that perceptual choices are biased towards the previous choice, a modulation that grows with previous decision confidence and previous response times, and are biased away from the direction of previous evidence, a modulation that grows with previous evidence strength. These findings suggest that previous choices and previous stimuli induce separate biases on subsequence choices through distinguishable and complementary mechanisms, pointing towards a complex process of decision formation.

## ACKNOWLEDGMENTS

This work was supported by a grant from the European Union Horizon 2020 Program (ERC Starting Grant 678286, “Contextvision”).

## Commercial relationships

none.

## Author contributions

EB, MF, and FPdL conceived and designed research; EB and MF developed experimental code; EB performed experiments; EB and BVE analyzed data; EB, MF, BE and FPdL interpreted results of experiments; EB and BVE prepared figures; EB drafted manuscript; MF, BVE, and FPdL edited and revised manuscript; EB, MF, BVE, and FPdL approved final version of manuscript.

